# Interplay of host regulatory network on SARS-CoV-2 binding and replication machinery

**DOI:** 10.1101/2020.04.20.050138

**Authors:** Shiek SSJ Ahmed, Prabu Paramasivam, Kamal Raj, Vishal Kumar, Ram murugesan, Ramakrishnan

## Abstract

We dissect the mechanism of SARS-CoV-2 in human lung host from the initial phase of receptor binding to viral replication machinery. We constructed two independent lung protein interactome to reveal the signaling process on receptor activation and host protein hijacking machinery in the pathogenesis of virus. Further, we test the functional role of the hubs derived from both interactome. Most hubs proteins were differentially regulated on SARS-CoV-2 infection. Also, the proteins of viral replication hubs were related with cardiovascular disease, diabetes and hypertension confirming the vulnerability and severity of infection in the risk individual. Additionally, the hub proteins were closely linked with other viral infection, including MERS and HCoVs which suggest similar infection pattern in SARS-CoV-2. We identified five interconnecting cascades between hubs of both networks that show the preparation of optimal environment in the host for viral replication process upon receptor attachment. Interestingly, we propose that seven potential miRNAs, targeting the intermediate phase that connects receptor and viral replication process a better choice as a drug for SARS-CoV-2.

## Introduction

Coronavirus disease 2019 (COVID-19, SARS-CoV-2) became a pandemic, spread almost 150 countries worldwide [1]. SARS-CoV-2 causes lower respiratory infections leading to pneumonitis and multi-organ failure [2]. Till date, 1991562 cases were reported as on 16th April 2020 and expected to increase every day by a human-to-human transmission that occurred due to close contact with one another [3]. An increase in patient number increases the fatality rate because of non-specific treatment against SARS-CoV-2. Today COVID-19 became a serious threat in both developed and under-developing countries [4]. There is an urgent need to accelerate work on specific treatments to decrease disease, morbidity and mortality [5, 6]. Repurposing of existing drugs has been considered as an immediate remedy rather than waiting for a new molecule to pass through human clinical trials. In addition, SARS-CoV-2 screening at the early stage may benefit the vulnerable population with the risk factor of diabetes, hypertension [7-9], and cardiovascular disease [10]. However, understanding the mechanism of SARS-CoV-2 and host involvement will provide knowledge that guides towards developing biomarkers and drugs to treat this pandemic disease.

SARS-CoV-2 is a single-stranded RNA virus-derived from the Human SARS-CoV-2 group of ß-genus with the genome size of 30kbp. SARS-CoV-2 genome encodes most structural proteins at 3’ ORFs and non-structural proteins (nsp) at 5’ end ORFs [8, 11]. The nsp proteins are categorized as nsp1-16, which plays a potential role in replicase / transcriptase activity whereas the ORFs at 3’ encode nine putative accessory factors along with four structural proteins, nucleocapsid (N), envelope (E), membrane (M), and spike (S) proteins [8, 11]. The envelope (E), membrane (M), and spike (S) proteins localized at vial surface suggested having host binding capacity. On host attachment, SARS-CoV-2 initiates the infectious cycle and undergoes RNA replication inside the host and assembles their substrate into their progeny virus. In general at the initial phase of the host binding, virus may create an optimal environment for its replication by interacting with the host proteins. Simultaneously, the host alters its gene expression to react against the virus for self-defense leading to the overexpression of inflammatory genes and cytokine storm [12]. Due to the viral genome complexity and 20% sequence dissimilarity with other pathogenic HCoVs [13]. Notably, the sequence variations at replicase complex (ORF1ab), envelope (E), spike (S), nucleocapsid (N) and membrane (M) suggest the possibility of different mechanisms adopted by SARS-CoV-2 that brings the question of repurposing the existing drugs. The current knowledge of host molecular components utilization by the SARS-CoV-2 is still at an early stage compared with other known human infecting RNA viruses. Understanding the regulatory behavior will enhance our knowledge of the virus-host mechanism, which may through light on diagnosis and treatment.

In this study, we first time report a systems biological framework (Fig 1) to investigate the molecular interplay between SARS-CoV2 and human lung tissue. The novel approach uses high through-put experimental data and human protein interactome to reveal the SARS-CoV2 driven host mechanism. Our approach provides (i) Functional hubs that activated upon viral attachment to the receptor and viral genome evasion to the host. (ii) Association of viral modulated functional hubs with diabetes, hypertension, and cardiovascular disease. (iii) An interdependency between the functional hubs that demonstrates the utilization of a host environment created upon receptor activation for the viral replication process. (iv) Hubs representing miRNA for therapeutic intervention. Overall, our framework accesses the information such as human tissue proteome, transcriptome, text mining, and miRNA data to postulate a mechanism in host on SARS-CoV2 infection.

**Fig 1.**
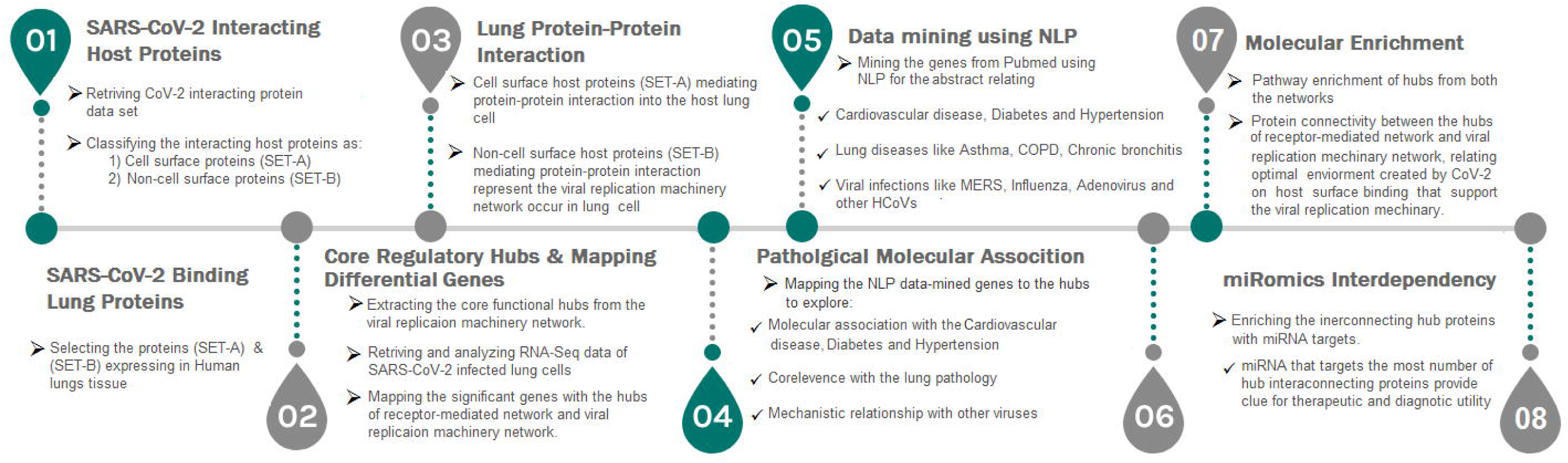
A strategic workflow diagram representing the step-wise processes of this study. Eight interdependent steps were followed, that majorly involved collection, classification, mapping, and data analysis. Each step was self-explained with the legend describing the process within the workflow.

## Results

### SARS CoV-2 Interacting host proteins

Firstly, we analyzed whether SARS CoV-2 protein interacting with the endogenous human proteins (host). For which we searched and collected SARS-Cov-2 host proteomics data derived from affinity purification mass spectrometry [14]. The data provide 332 human endogenous proteins interacting with 27 SARS CoV-2 proteins. Further, these human endogenous proteins were categorized as SET-A and SET-B, respectively. SET-A composed of 38 human proteins showing physical affinity with structural proteins like as envelope (E), spike (S), and membrane (M) proteins localized at SARS-CoV2 surface. Whereas, the SET-B contains 294 host proteins confirming its affinity with 23 SARS-CoV2 non-structural and one nucleocapsid (N) proteins. Then, we investigate host proteins that favor the attachment of SARS-CoV2. So, the cellular localization of SET-A proteins was annotated to have ATP6V1A, AP3B1, STOM, and ZDHHC5, localized at the host cell surface that may enable SARS-CoV2 attachment. Further, the proteins in the SET-A and B were mapped with an in-house *lung-expressed gene* database to confirm their expression in the lungs that relate the mechanism that occurs in lung tissue.

### Lung Protein-Protein Interactions

Next, we investigate the high-through-put protein-protein interaction networks that reveal the SARS-CoV2 mechanism in the host. Two independent protein interactome networks were generated from SET-A and SET-B, respectively. The first interactome, receptor-mediated network from SET-A contains four cell surface seed proteins that extended to have 45 neighboring proteins with every node directly connected to its seed proteins. The receptor-mediated network describes the molecular signaling initiated on SARS-CoV2 attach to the host. Secondly, the viral replication machinery network from SET-B with 332 seed proteins extended to 1486 neighboring proteins with 11438 interacting edges which representing the mechanism attributed to evasion of the SARS-CoV2 genome into the host. The proteins involved in the networks were mapped to the in-house *lung-expressed-gene* database. As a result, all 49 proteins with four seed in the receptor-mediated network were expressed in the lungs (Fig 2). Similarly, the viral replication machinery network was acquired with 1522 proteins with 9747 interacting edges showing the complex SARS-CoV-2 mechanism in the human lungs.

**Fig 2.**
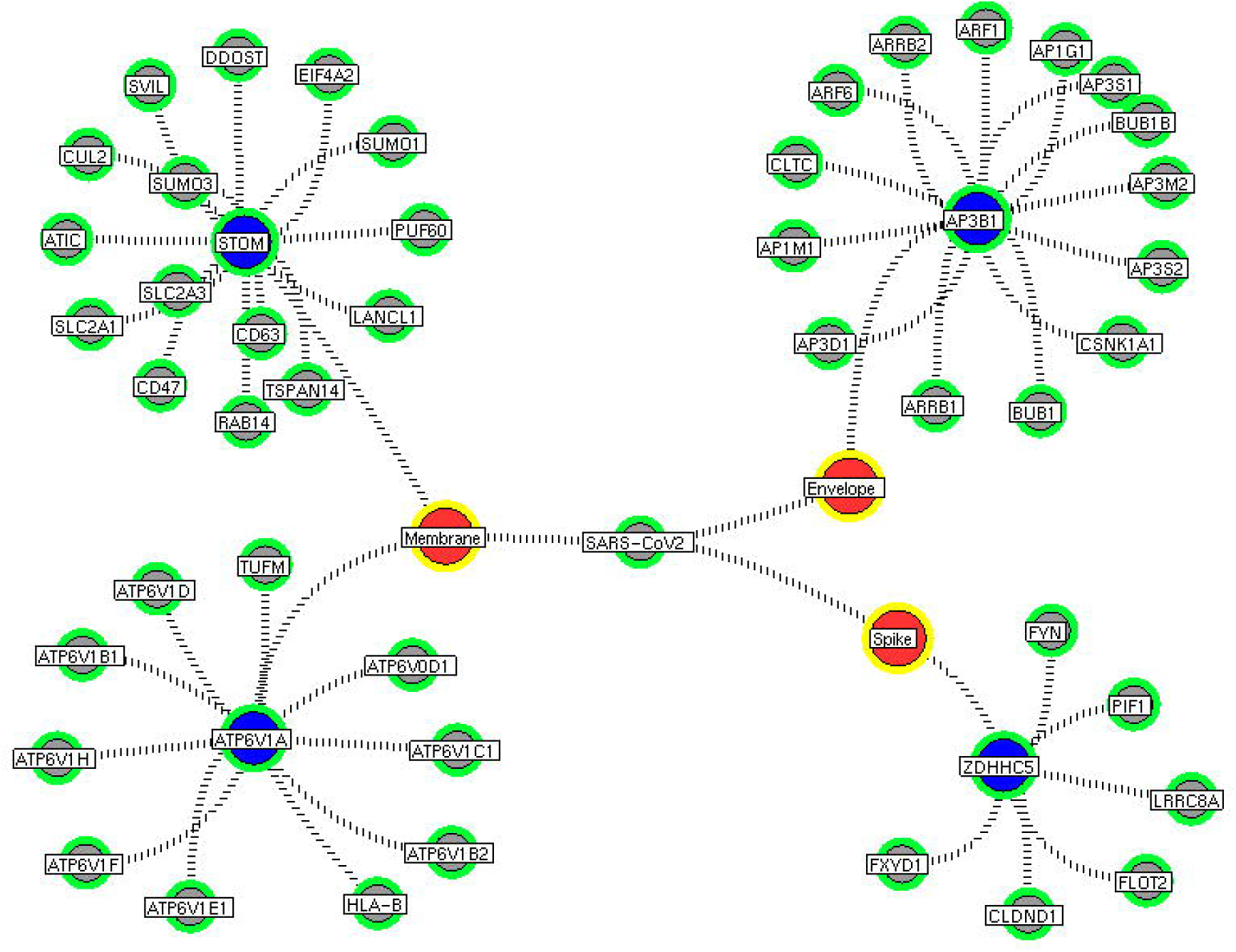
Receptor-mediated network. The protein interaction network describing the protein connectivity between the host and SARS CoV2. The three red color nodes represent the SARS CoV2 surface proteins interacting (black dotted) with the seed proteins, STOM, ATP6V1A, AP3B1, and ZDHNNC5, a host protein (violet) localized at the cell surface. All four seed proteins mediated it signals to its interacting proteins (green) and form a four individual closed hub.

### Core regulatory hubs

Here, we looked for densely connected protein, which provides essential functional hubs within the network. Molecular complex detection (MCODE) algorithm [15] used to extract hubs that define the core regulatory signaling and cellular processes within the network. Eleven core functional hubs were obtained from the viral replication machinery network (S1-11 Fig). In a receptor-mediated network, the implementation of the MCODE algorithm was exempted for its limited number of nodes and edges, which by itself forms hubs. Hence, the interacting proteins for all four distinct receptors were considered as hubs for the receptor-mediated network (Fig 2). All hub proteins were mapped to differential expressed genes from RNA-Seq data. Of 374 hub proteins from both the network, 76 were differentially regulated in primary human lung epithelium (NHBE) on SARS-CoV2 infection (RNA-Seq dataset GSE147507). For instance, eight differentially expressed hub proteins were noticed in the receptor-mediated network (S1 Table). Similarly, among 325 hub proteins of viral replication machinery network, 68 were altered significantly, which confirms the hijacking of endogenous human proteins for the viral replication process (S1 Table).

### Data Mining and Molecular risk factor Association

Next, we investigated why SARS-CoV2 infection is susceptible to metabolic diseases like cardiovascular disease, diabetes, and hypertension. For that, we tested the associations of hub proteins with these metabolic diseases using natural language processing in our indigenous R-code. Among 374 hub proteins from the network, 186, 155, and 103 proteins were associated with cardiovascular disease, diabetes, and hypertension, respectively (S2 Table). We adopted a similar mining process to determine the relevance of hub proteins with lung diseases such as asthma, chronic bronchitis, pneumonia, chronic obstructive pulmonary disease (COPD), and emphysema. Simultaneously, the relevance of hub proteins was tested in viral infections such as adenovirus, influenza, metapneumo, parainfluenza, respiratory syntical virus, rhinovirus, HCoV-NL63, HCoV-229E, HCoV-HKU1, HCoV-OC43, SARS-CoV and MERS-CoV. These results suggest that 101 hub proteins have relevance to lung disease (S3 Table), and 117 proteins were associated with the known viral infection (S4 Table). These associating hubs convey the likelihood of vulnerability in the risk population, symptoms related to lung diseases, and similar modes of viral infection.

### Molecular Enrichment

Subsequently, the protein enrichment analysis was executed to check the molecular involvement of hubs on SARS-CoV2 infection. The enrichment analysis of hubs showed several over representing pathways for the hubs of receptor-mediated (Fig 3) and viral replication machinery network (Fig 4). The receptor-mediated network hubs exhibit several molecular signaling and cellular events upon SARS-CoV2 attachment (S5 Table). Notably, most hub proteins showed significant association with immune mechanism, signal transduction, metabolism of proteins, metabolism of RNA, transcription machinery, vesicle-mediated transport, metabolism, transport of small molecules cell cycle and homeostasis were noticed (S5 Table). Also, a moderate involvement was seen in cellular response mechanism, cell-cell communication, organelle biogenesis and maintenance. Similarly, the enrichment analysis of hubs derived from viral replication machinery network (Fig 4) showed a major involvement of immune mechanism, signal transduction, metabolism of proteins, metabolism of RNA, transcription machinery, DNA repair, DNA replication, programmed cell death, and cellular responses to external stimuli were noticed (Table S5). Interestingly, several common mechanisms were noticed that occur due to the existence of 34 interconnected proteins between the hubs of both the network (Fig 5-9). These common molecules represent the inter-connecting mechanism involved in the transcription machinery, immune response, cell growth and/or maintenance, transport, metabolism, protein metabolism, cell communication and signal transduction that activated upon virus binding and has been subsequently utilized for viral replication process (S5 Table) (Fig 10).

**Fig 3.**
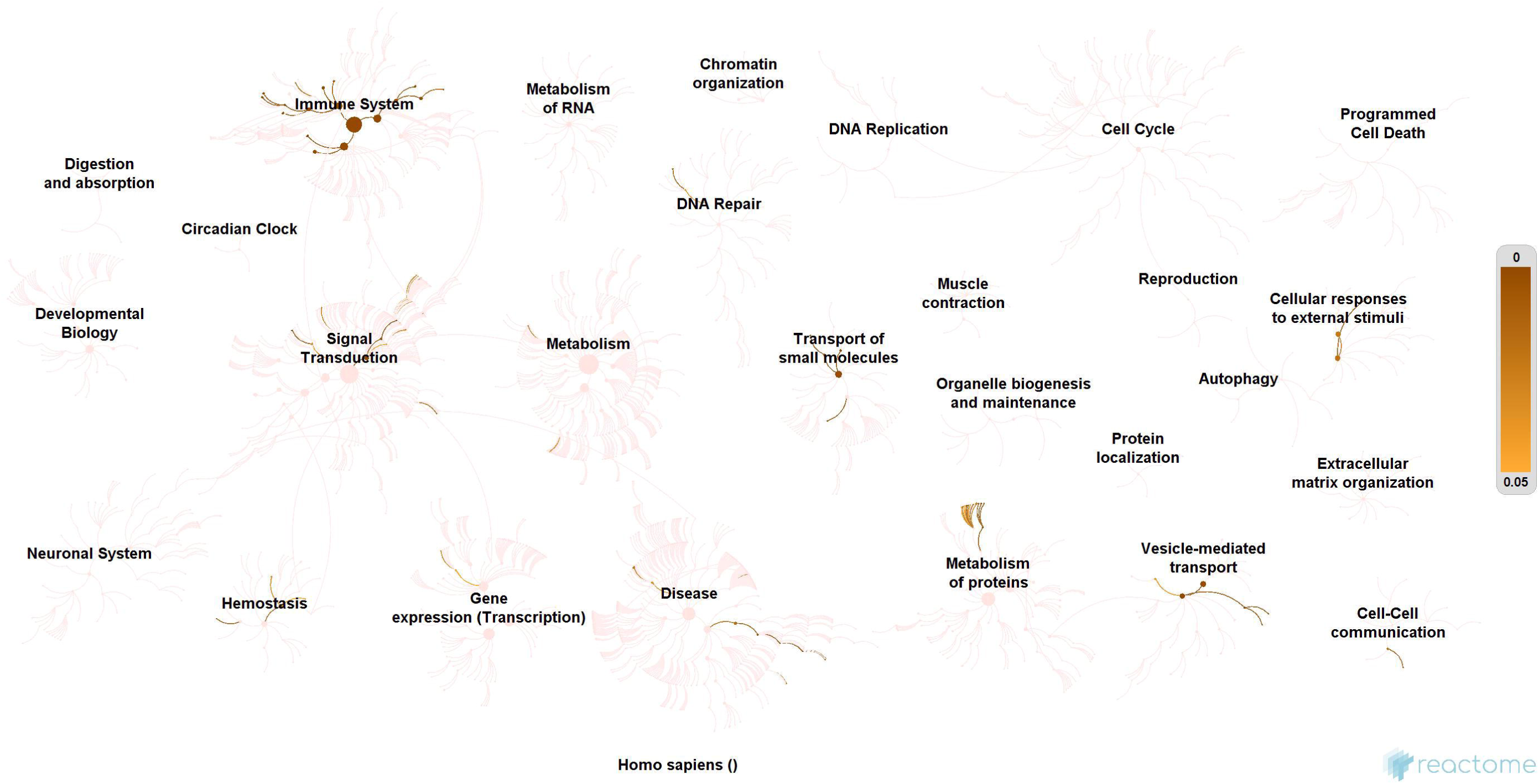
Molecular enrichment of receptor hubs. Enrichment of hubs showing major involvement of immune system, metabolism, and signal transduction. Significant involvement of hemostasis, transcription machinery, transport of small molecules, cellular response to external stimuli, protein metabolism, and vesicle-mediated transport. Over-representation of these molecular events suggests host self-defense mechanism and preparation of the optimal environment in the host for the viral replication process on receptor attachment.

**Fig 4.**
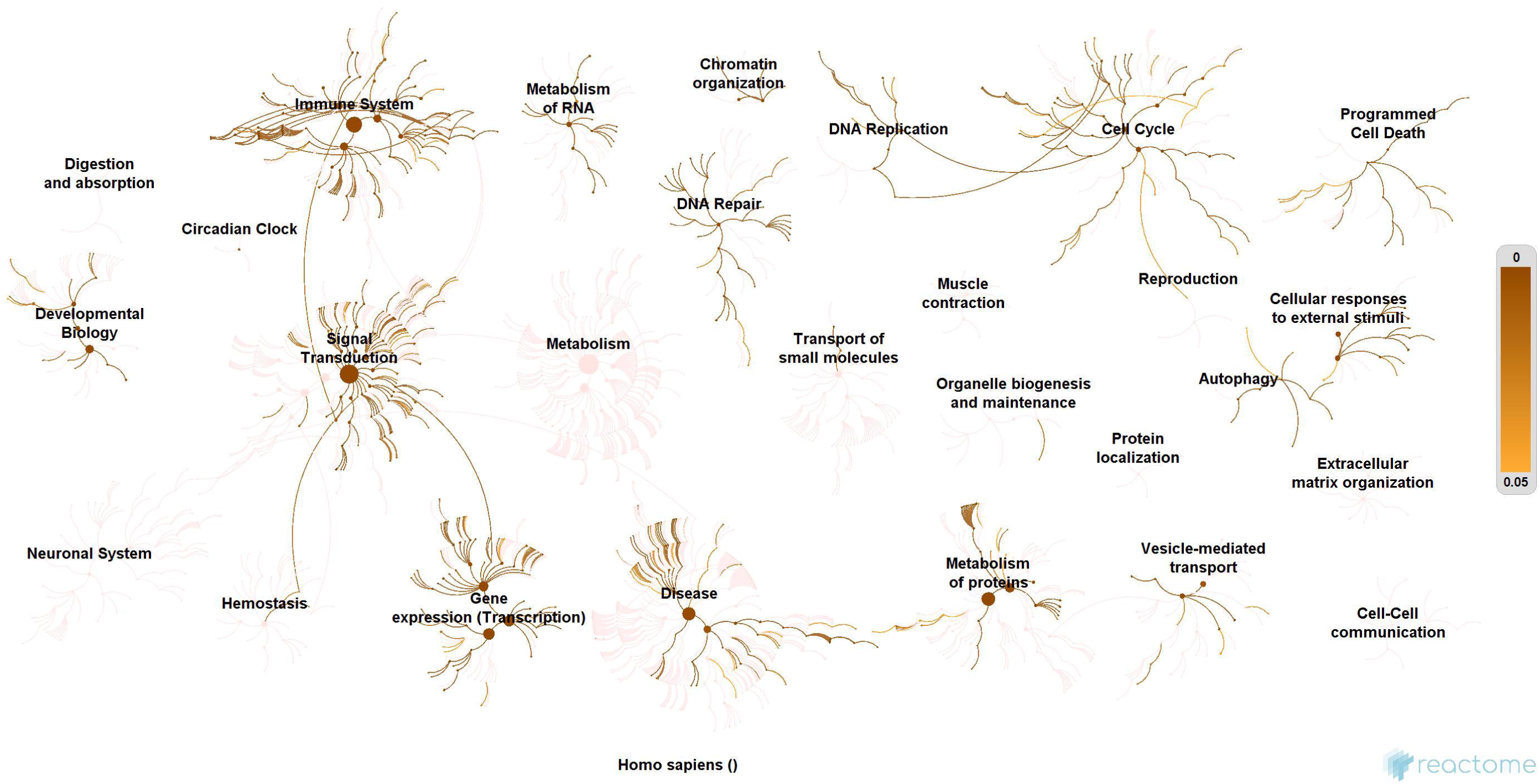
Enrichment of hubs derived from virus replication machinery network. Enrichment of 325 proteins in eleven hubs showing the wide coverage of involvement in the immune system, signal transduction and active involvement of transcription machinery, protein metabolism, cell cycle, metabolism, DNA replication, DNA repair, metabolism, programmed cell death, and cellular response to external stimuli.

**Fig 5.**
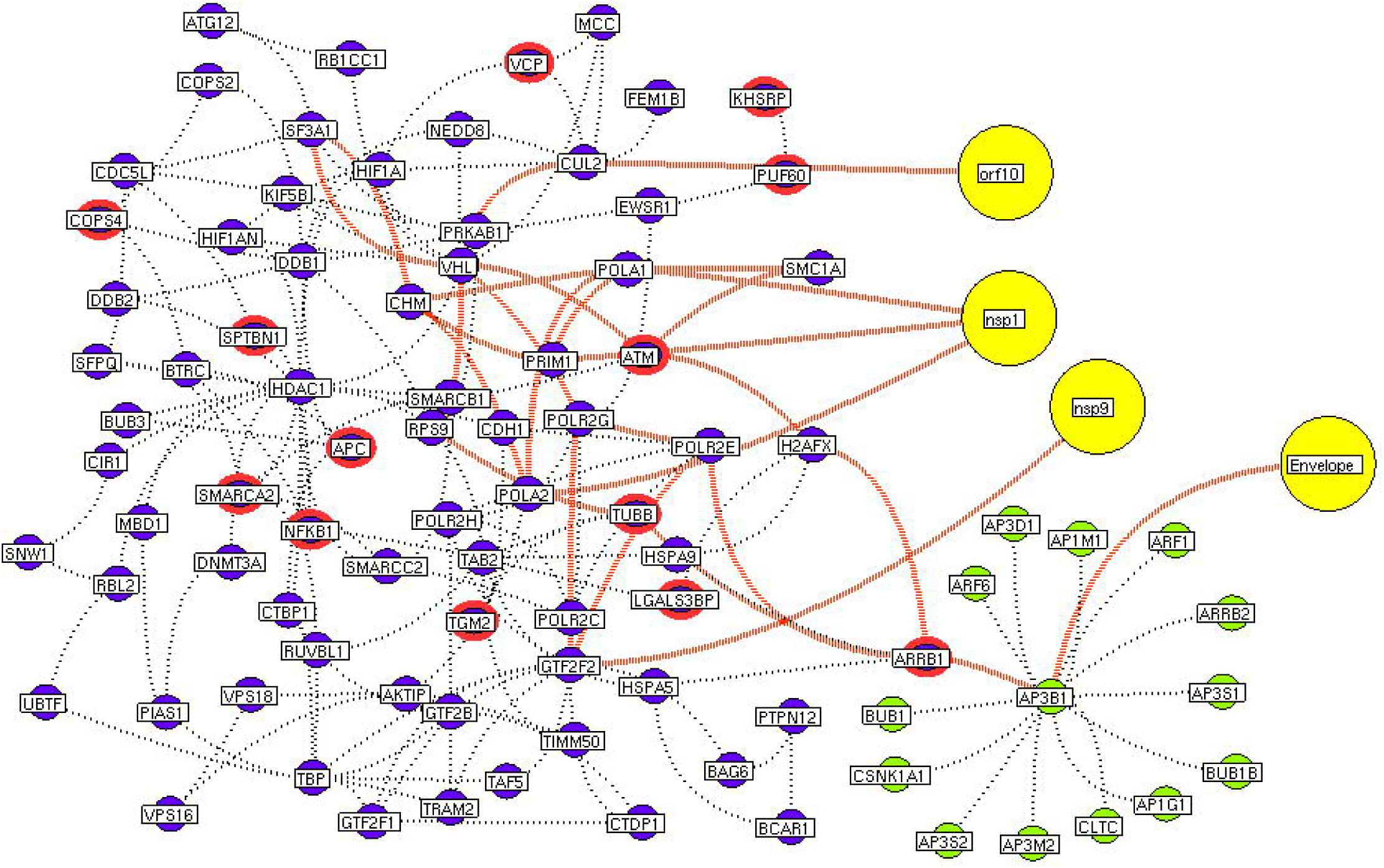
Protein interconnectivity between the functional hubs. The hub network of AP3B1 hub (green node) and hub 4 (violet node) demonstrating the interconnecting proteins between (red dotted edges) the hubs. The yellow nodes represent the viral protein interaction with the hubs. The viral envelope interacts with host AP3B1 surface protein that activates a series of proteins leading to utilization for viral machinery propose of orf10, nsp1 and 9. Additionally, the red highlighting node represents the differentially expressed genes on SARS-CoV2 infection derived from RNA-Seq analysis.

**Fig 6.**
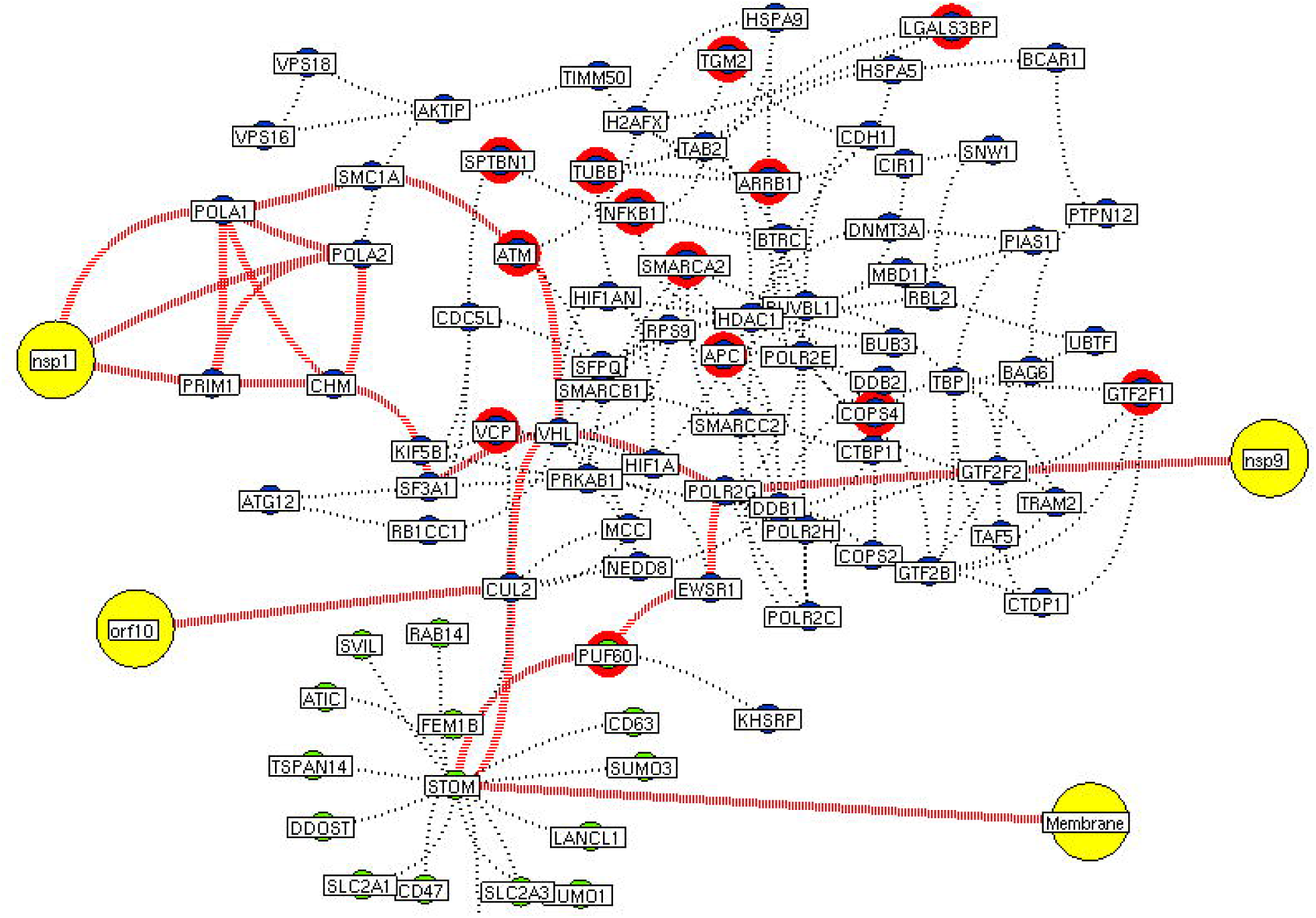
Interconnectivity between the functional hubs. The hub network of STOM hub (green node) and hub 4 (violet node) demonstrating the interconnecting proteins between (red dotted edges) the hubs. The yellow nodes represent the viral protein interaction with the hubs. The viral membrane interacts with host STOM surface protein that activates the series of proteins leading to utilization for viral machinery propose of orf10, nsp1, and 9. Additionally, the red highlighting node represents the differentially expressed genes on SARS-CoV2 infection derived from RNA-Seq analysis.

**Fig 7.**
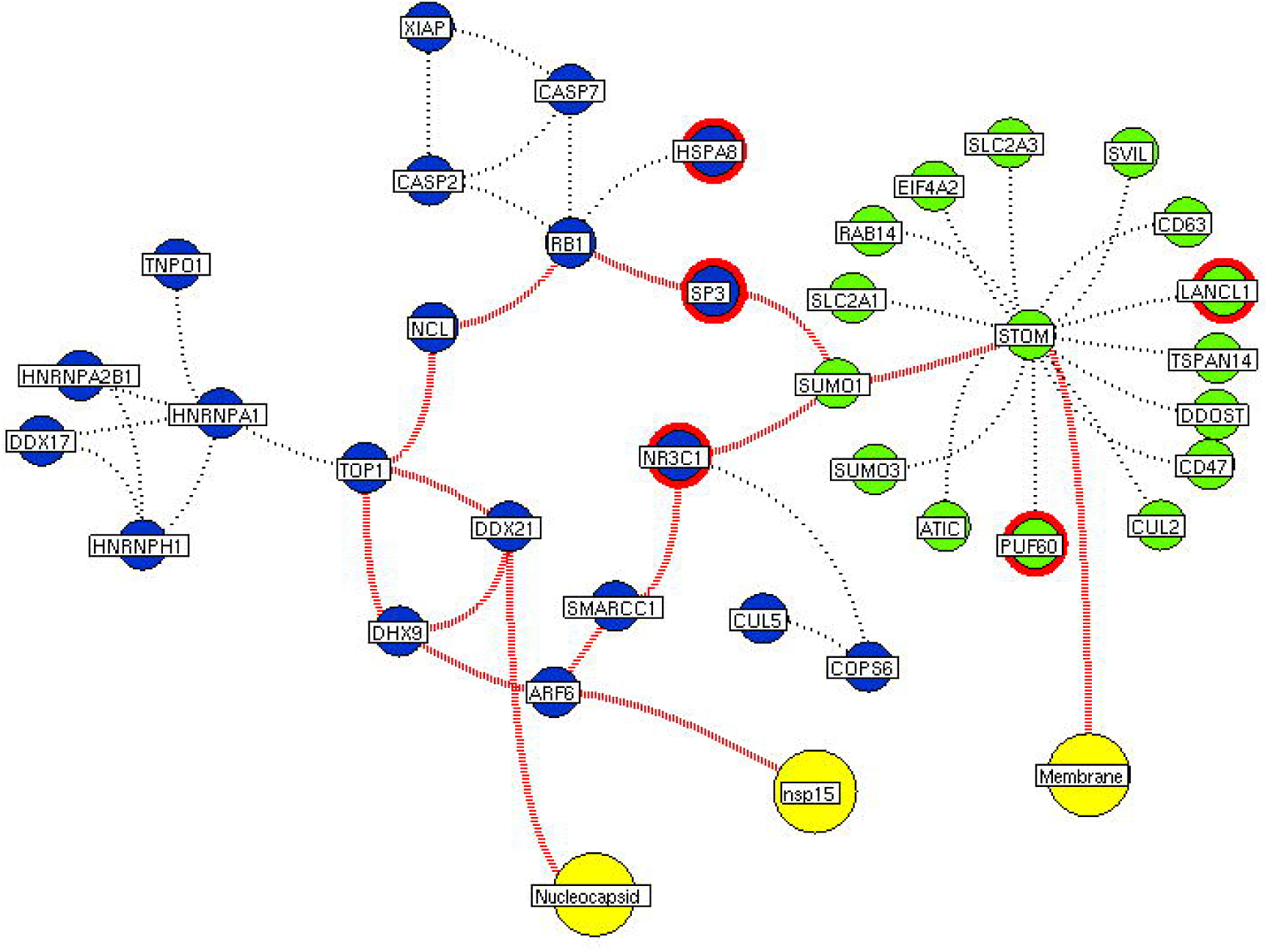
Interconnectivity between the functional hubs. The hub network of STOM hub (green node) and hub 7 (violet node) demonstrating the interconnecting proteins between (red dotted edges) the hubs. The yellow nodes represent the viral protein interacting with the hubs. The viral membrane interacts with host STOM surface protein that activates the series of proteins leading to benefit viral machinery relating to nucleocapsid and nsp15. Additionally, the red highlighting node represents the differentially expressed genes on SARS-CoV2 infection derived from RNA-Seq analysis.

### miRomics Interdependency

We further investigate these 34 interconnecting proteins that may use for a therapeutic process. For which, the miRNA target for interconnecting proteins (genes) was identified. Among 34 proteins, 26 shown targeted by 119 miRNA. Of which, miR-124-3p, let-7g-5p, miR-133a-3p, miR-133b, miR-218-5p, miR-22-3p and miR-506-3p were noticed to target more than three interconnected proteins (Fig 11). These miRNAs may show its utility for therapeutic intervention.

**Fig 8.**
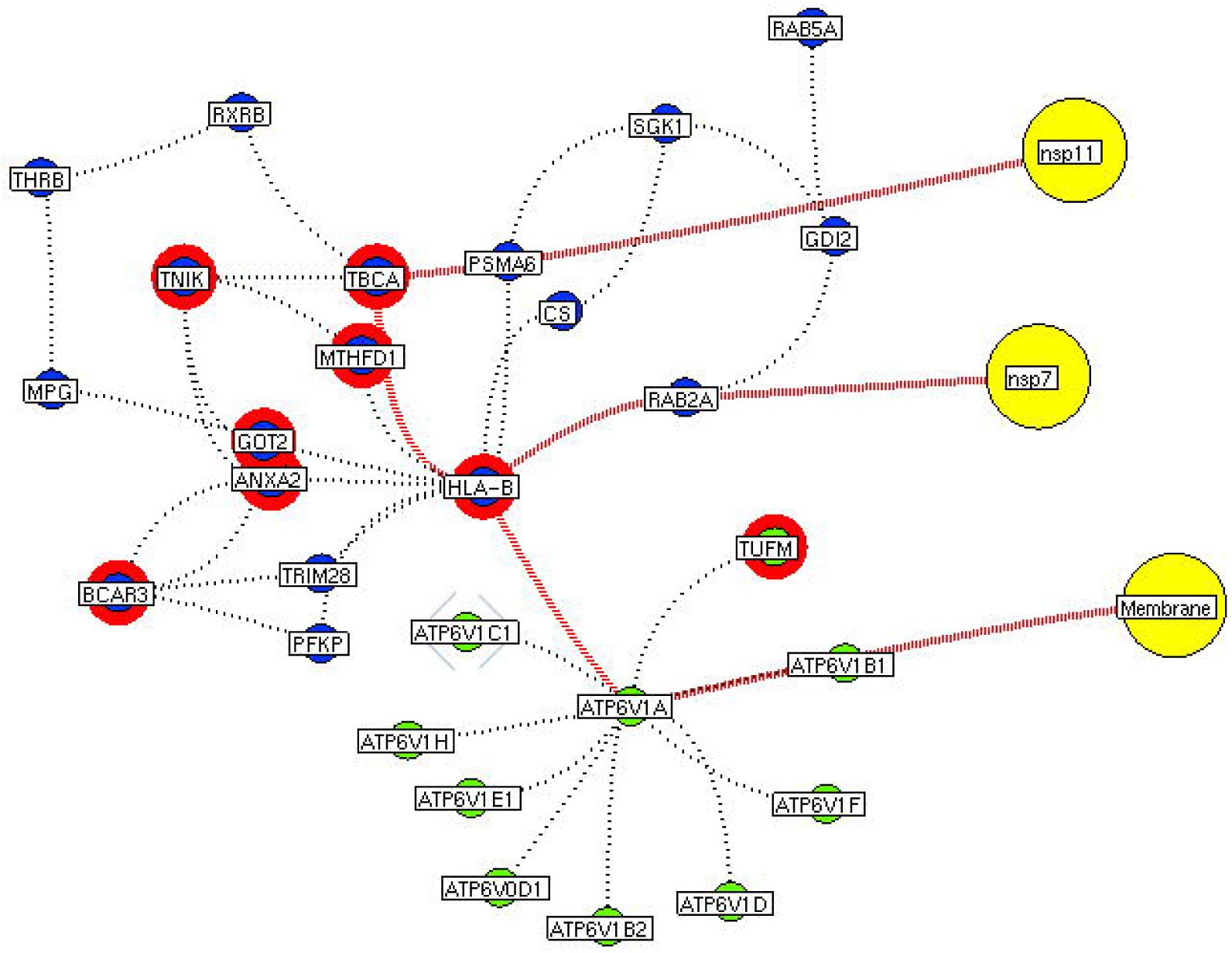
Functional hubs interconnectivity. The hub network of ATP6V1A hub (green node) and hub 5 (violet node) demonstrating the interconnecting proteins between (red dotted edges) the hubs. The yellow nodes represent the viral protein interacting with the hubs. The viral membrane interacts with host ATP6V1A surface protein that activates the series of proteins leading to benefit viral machinery relating to nsp7 and nsp11. The red highlighting node represents the differentially expressed genes on SARS-CoV2 infection derived from RNA-Seq analysis.

**Fig 9.**
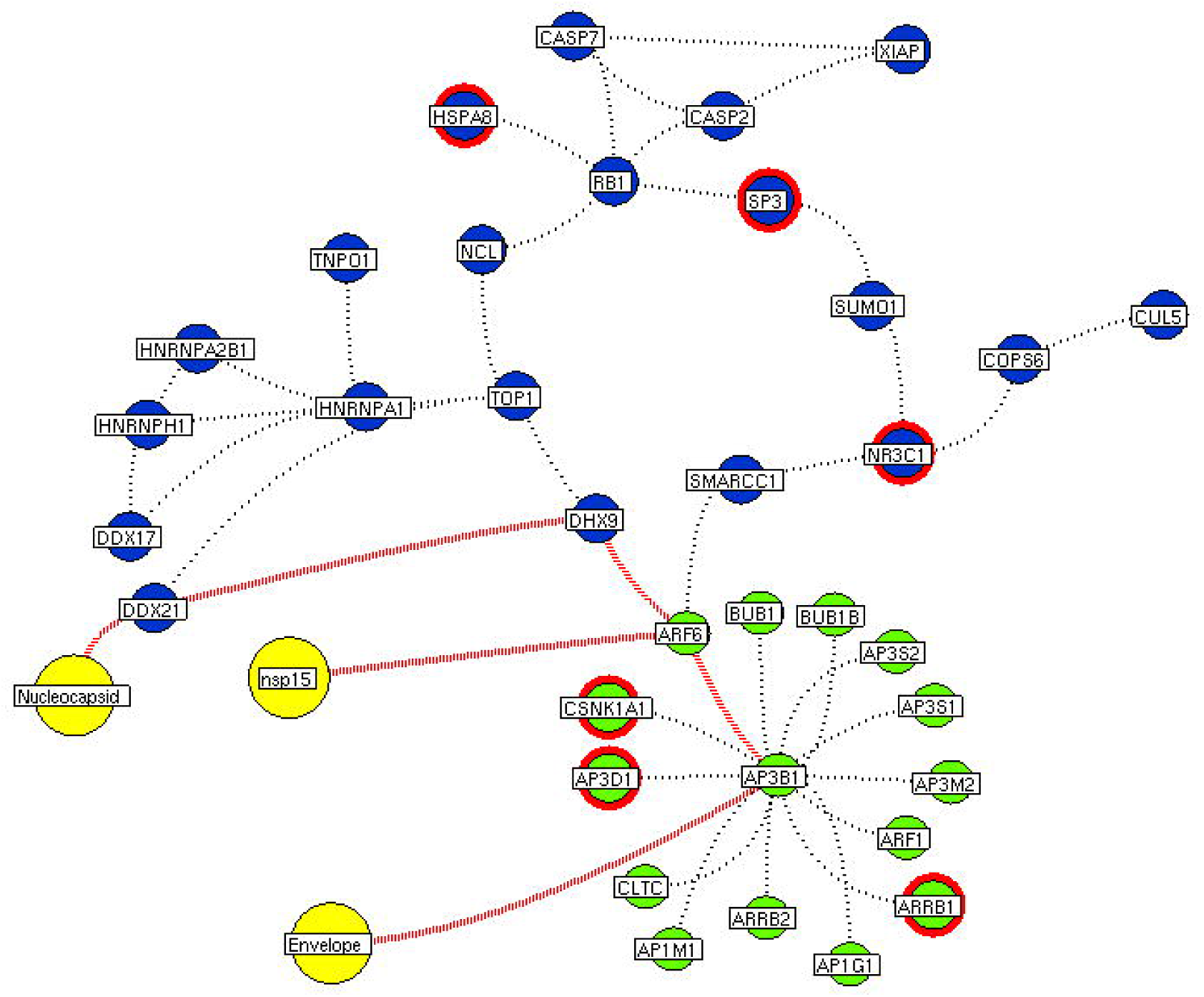
Functional hubs interconnectivity. The hub network of ATP6V1A hub (green node) and hub 5 (violet node) demonstrating the interconnecting proteins between (red dotted edges) the hubs. The yellow nodes represent the viral protein interacting with the hubs. The viral envelope interacts with host ATP6V1A surface protein that activates the series of proteins leading to benefit viral machinery relating to nucleocapsid and nsp15. Additionally, the red highlighting node represents the differentially expressed genes on SARS-CoV2 infection derived from RNA-Seq analysis.

**Fig 10.**
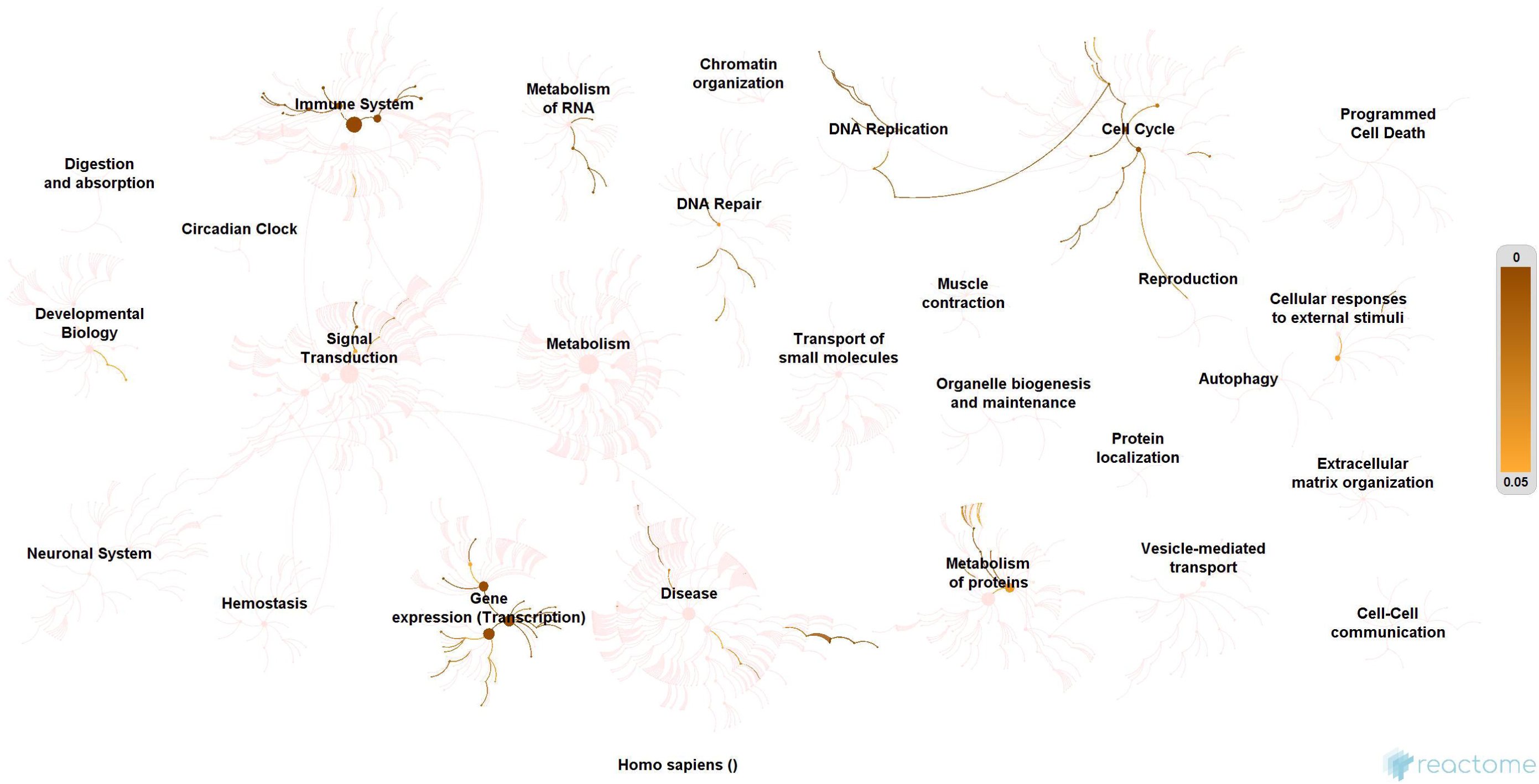
Enrichment of interconnecting hub proteins. Enrichment of 34 proteins from five interconnected hubs showing the contribution in the immune system, signal transduction, transcription machinery, and protein metabolism. These interconnecting mechanisms suggest being the intermediate phase that connects the events between early host attachment and the viral replication process.

**Fig 11.**
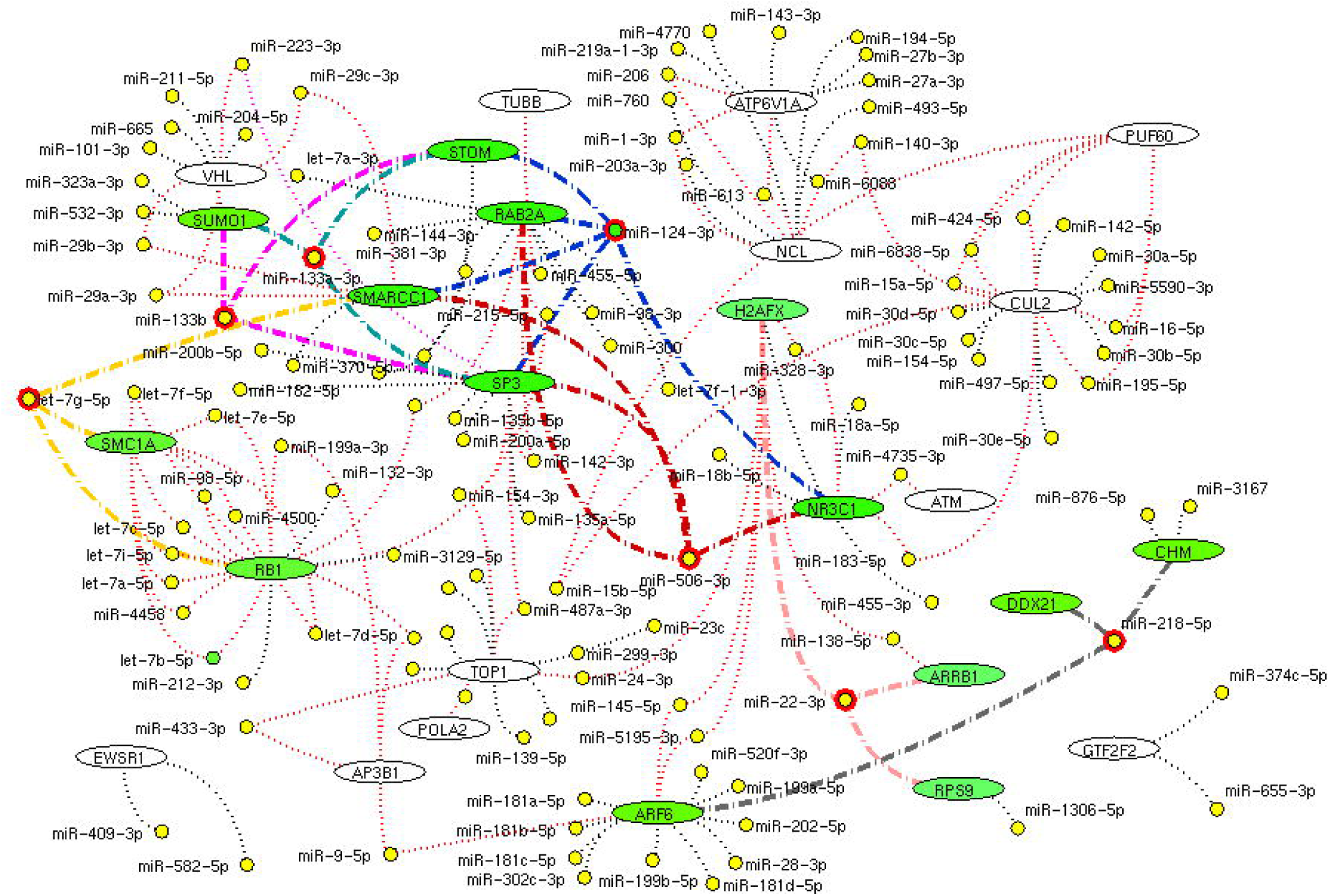
miRNA network. The network demonstrates the miRNA (yellow node) target for (black dotted edges) 34 protein encoding genes (while ellipse). The miRNA (red highlighted node) that inhibits (bold edges) more than three genes (green ellipse) interconnecting hub proteins are represented. These miRNAs will be the potential molecule as a drug that helps in inhibition of connective between host attachment and the viral replication process.

## Discussion

We implement a comprehensive systems biological framework (Fig.1) that utilizes a variety of datasets to illustrate the critical mechanism mediated by SARS-CoV2 in the lungs. Two independent lung proteins interactome were constructed. The first interactome termed as a receptor-mediated network that revealed the molecular interaction that mediates cellular and signaling processes on SARS-CoV2 attachment to the host. In the receptor-mediated network, ATP6V1A, AP3B1, STOM, and ZDHHC5 are the seed proteins localized at the host surface showed significant affinity to the outer surface envelope (E), spike (S), and membrane (M) proteins of SARS-CoV2. On bounding with ATP6V1A, AP3B1, STOM, and ZDHHC5, the host activates multiple signaling and cellular events that demonstrated in the receptor-mediated protein network (Fig 3). For instance, viral envelope (E) bound to the AP3B1 cell-surface protein that initiates the interaction with its neighboring proteins, AP1G1, AP1M1, AP3D1, AP3M2, AP3S1, AP3S2, ARF1, ARF6, ARRB1, ARRB2, BUB1, BUB1B, CLTC, and CSNK1A1. Similarly, spike (S) interacts with the ZDHHC5 that connects CLDND1, FLOT2, FXYD1, FYN, LRRC8A, and PIF1. Likewise, the SARS-CoV2 membrane (M) interacts with ATP6V1A and STOM host surface protein that signals 25 human endogenous proteins (Fig 2). Each host cell surface proteins and it’s interacting neighbors form a closed hub with no extensive interconnection with another hub. These hubs were pathways enriched that takes part in wide verities of molecular pathways that include endosome transport, vesicle-mediated transport, protein modification, regulation of cell cycle, immune response, kinase activity, signal transduction, protein metabolism, cell communication, energy and metabolic pathway, cell growth, and maintenance, regulation of nucleic acid metabolism. To our knowledge, most of this mechanism is very well connected with the viral pathogenesis machinery mechanism. Notably, at an early phase of surface attachment, the host initiates inflammatory signaling such as interleukin signaling (IL1, IL2, IL5, IL23, IL8), TNF receptor, and NF-kappa B signaling pathways that may initiate cytokine storm in lung cells [12, 16]. Simultaneously, the involvement of transcription and translational regulatory processes were noticed in the pathway enrichment of receptor hubs. These results suggest that on surface receptor activation host prepares an optimal environment in the host for the viral replication. The second lung protein interactome was named as viral replication machinery network. In this network, we demonstrated the complex interconnectivity of human endogenous proteome with nucleocapsid (N) and 23 non-structural SARS-CoV2 proteins. The viral replication machinery network conveys the utilization of host factors for SARS-CoV2 replication and self-defense mechanism. Using the MCODE algorithm, eleven hubs (S1-11Fig) were extracted from the viral replication machinery networks, which play a significant role in cellular and molecular mechanisms (S5 Table). Further, mapping the differently expressed genes with the hubs showed approximately 20% of hub proteins were altered upon SARS-CoV2 infection (S1 Table). This result confirms the hijack of host hub proteins and their molecular pathways for SARS-CoV2 machinery. Some hub proteins were proven linked to diabetes, hypertension, and cardiovascular disease (S2 Table). These results suggest that the SARS-Cov-2 alter and utilize the disease proteins for its replication machinery that may be the considerable cause for susceptibility, severity, and complication. Besides, similar observations were noticed while mapping hub proteins with proteins related to lung disease and other viral infections. On mapping, 101 hub proteins were shared between the lung diseases suggesting a common mechanism relating disease symptoms and/or vulnerability (S3 Table). Also mapping with other viral infection dataset, 50 hub proteins of the replication machinery network have noticed in influenza virus infection (S4 Table), which suggests SARS-CoV2 and influenza may have a similar mode of host infection machinery [17].

As apart, several common pathways between the hubs of two networks were noticed (S5 Table). We obtained these common pathways as an intersection of the 34 interacting molecules between the hubs (Fig 5-9). For example, AP3B1 hub derived from the receptor-mediated network interact with the “hub 4” of viral replication machinery network by GTF2F2, POLR2E, ARRB1, and AP3B1 interconnecting proteins (Fig 5). Overall, five interacting hubs were noticed between receptor-mediated and viral replication machinery network (Fig 5-9). The molecular pathways of interconnecting protein hubs could be the intermediate phase that connects the receptor activation mechanism and viral replication process (Fig 10). Such intersecting proteins will aid in developing a drug for SARS-CoV2.

Here, we suggest a few miRNAs that regulates the proteins of interconnecting hubs, which may have therapeutic potential. The miRNA has been regarded as drug molecules for various diseases [18 19]. Of 111 miRNA, miR-124-3p, let-7g-5p, miR-133a-3p, miR-133b, miR-218-5p, miR-22-3p and miR-506-3p targets minimum of three interconnecting proteins (Fig 11). These seven miRNA molecules are needed to be viewed in a better perspective for future research to screen and treat the current SARS-CoV2 pandemic. Overall, this study displays several advancement and advantages, 1) Establishes interplay between SARS-CoV2 and human lung host mechanism by integrating experimental evidence arrived from high throughput multi-omics observation on SARS-CoV2. 2) Our approach provides a possible clue for susceptibility and severity in an individual with metabolic disease complications and suggests possible proteomic relevance with lung diseases and other viral infections. 3) Our study increases knowledge of molecular interconnectivity between receptor binding mechanism and viral replication machinery that guide towards drug target research. Although our approach provides the multi-dimensional view on SARS-CoV2 and host interaction, two major limitations need to be considered, 1) the interaction between SARS-CoV2 spike protein with the host angiotensin (ACE2) receptor is not defined in affinity purification mass spectrometry data. Hence, no direct ACE2 interaction has been established in our interactome. 2) We propose a few miRNAs from our approach, are needed to be validated further for clinical application.

## Conclusion

In summary, our systems biological approach provides an extensive investigation on SARS-CoV2 host interaction by constructing interactome based on experimental evidence. We identified crucial functional hubs that relate the mechanism on activation SARS-CoV2 attachment through receptor and subsequent utilization functional hubs for viral replication in the host. Enrichment analysis supports our hypothesis showed the common mechanism associated with transcriptional and protein translation processing activated upon in host attachment. Hub proteins showed linked with diabetes, hypertension, and cardiovascular disease proteins that provide the clue for severity and susceptibility in the diseased population. The relationship between hub proteins with lung diseases like COPD, asthma, pneumonia establishes the likelihood of similar symptoms. Also, assessing the hubs with the reported proteins of other viral infection establishes the similarity in the mode of viral infection. Interestingly, we propose miR-124-3p, let-7g-5p, miR-133a-3p, miR-133b, miR-218-5p, miR-22-3p, and miR-506-3p from the genes of interconnecting proteins from its molecular pathways as a miRNA drug for SARS-CoV2. However, further work is needed to confirm its utility for clinical application. We believe our results will add knowledge to the existing information that may open up miRNA as a drug for SARS-CoV2 infection.

## Methods

### SARS CoV-2 Interacting host proteins

The dataset relating severe acute respiratory syndrome coronavirus 2 (SARS-CoV2) and human proteins interactions searched electronically. Of extensive search, the experimentally proven dataset reporting the physical association of 26 SARS-CoV2 proteins with human proteins determined using affinity purification mass spectrometry (AP-MS) [14] was retrieved. We converted all collected interacting human proteins to an official gene/protein symbol using the Uniport database (https://www.uniprot.org/). Further, the interacting proteins were categorized into SET-A and SET-B. The SET-A contains Human proteins that interact with SARS-CoV2 structural proteins, especially envelope (E), spike (S), and membrane (M) whereas SET-B contains Human proteins that interact with nucleocapsid (N) and 22 non-structural proteins. The SET-A was further curated to have proteins localized at plasma membrane using Cell surface protein atlas (https://wlab.ethz.ch/cspa/), Human Protein Reference Database (HPRD) (https://www.hprd.org/), and Human Protein Atlas (https://www.proteinatlas.org/) database, assuming SARS-CoV2 structural proteins use these host surface proteins for attachment.

### SARS CoV-2 binding human lung proteins

Next, we collected lung protein expression data from the HPRD (https://www.hprd.org/), Human Proteome Map (http://www.humanproteomemap.org/), Proteomics DB (https://www.proteomicsdb.org/) and Human Protein Atlas (https://www.proteinatlas.org/) database and integrated using Microsoft access based on the common identifier to create an in-house *lung-expressed-gene* database. Each dataset (SET-A and B) was mapped to the *lung-expressed-gene* database to confirm all regulatory and binding events occur in Human lung tissue.

### Lung Protein-Protein Interactions

For SET-A and B, we collected human protein-protein interaction data using HumanProteinpedia (http://www.humanproteinpedia.org/), BioGRID (https://thebiogrid.org/), HPRD (https://www.hprd.org/), and Intact (https://www.ebi.ac.uk/intact/) database. All collected interactions were mapped to the *lung-expressed-gene* database to construct human lung protein-protein interactome. Each protein interaction in SET-A and SET-B was considered if both the interacting proteins neighbors were expressed in lung tissue. Thereafter, SET-A was designated as a receptor-mediated network and SET-B was termed as a viral replication machinery network.

### Core regulatory hubs

MCODE algorithm [15] was executed on the viral replication machinery network to decompose the large network into dense functional hubs with a minimum of three proteins in each hub. The derived functional hubs from the viral replication machinery network describe the core functional module of host influenced by the non-structural proteins of SARS-CoV2. Whereas in the receptor-mediated network, implementing MCODE algorithm was exempted because of its limited number of nodes and edges, which by itself forms hubs. The functional hubs of the receptor-mediated network exhibit the mechanism that activated upon SARS-CoV2 attachment to the lung surface proteins. Subsequently, in order to determine the involvement of each hub in viral pathogenesis, the expression data of SARS-CoV2 infection was analyzed, and the differential expressed genes were mapped hub proteins.

### Mapping the Differential Genes

We searched the gene expression dataset in NCBI Gene Expression Omnibus GEO (http://www.ncbi.nlm.nih.gov/geo/) using various key terms related to SARS-CoV2. Our search showed the availability of RNA-Seq dataset (GSE147507) executed in primary human lung epithelium (NHBE) infected with SARS-CoV2 (https://www.ncbi.nlm.nih.gov/geo/query/acc.cgi?acc=GSE147507). The raw FASTQ files (SRX7990866) of NHBE infected with SARS-CoV2 and their respective controls were retried from NCBI, SRA database. After quality assessment, the reads were aligned to the human genome (hg19) following the RNA-Seq analysis pipeline to determine the differential expression genes using DESeq with p-value < 0.05. Further, the significant differentially expressed genes in the host on SARS-CoV2 infection were collected and mapped to the hubs.

### Data Mining and Molecular risk factor Association

R-program was employed to find out the associated genetic hubs risk factors genes reported in the literature for diabetes, hypertension, and cardiovascular diseases. In brief, the R-code collects the abstract from the NCBI, PUBMED, for the keywords related to each risk factor. All abstracts were automatically examined for the presence of risk factors (examples: “diabetes”) and hub proteins using a natural language processing method. Further, the co-occurrence of risk factors with each protein symbol in the abstract was assessed using a point-wise mutual information method. A similar data mining process was carried out for lung diseases (asthma, chronic bronchitis, pneumonia, and emphysema) and other viral pathogens such as adenovirus, influenza, metapneumo, parainfluenza, respiratory syntical virus, rhinovirus, HCoV-NL63, HCoV-229E, HCoV-HKU1, HCoV-OC43, SARS-CoV and MERS-CoV.

### Molecular Enrichment

Simultaneously, all hubs were enriched using Kyoto Encyclopedia of Genes and Genomes (Reactome; https://reactome.org//) database to evaluate the relevance of the pathway of each hub in SARS-CoV2 infection and replication process. In particular, the pathway enrichment of receptor-mediated network hubs shows the mechanism initiated upon SARS-CoV2 receptor activation. Whereas, the pathways of viral replication machinery network hubs explain the mechanism attribute on evasion of SARS-CoV2 genome into the host for its replication process.

### miRomics Interdependency

miRomics enrichment analysis was performed for the selected genes in hubs considering the importance of miRNA in treatment. The relevant information gene-miRNA target was collected from the electronic database miRTarBase Version 7.0 [http://mirtarbase.mbc.nctu.edu.tw/php/index.php]. The most targeting miRNA to interconnecting proteins were identified and proposed as a possible drug for SARS-CoV2.

## Acknowledgments

All authors thank Chettinad Academy of Research and Education (CARE) for the support. We specially thank Professor Ram Murugesan, Research Director (CARE) for the critical comments and feedback on the manuscript.

## Supporting information

**S1 Fig. Functional hub derived from viral replication machinery network**. This cluster is termed as hub 1 that represents the closed interconnected proteins (rectangular box) showing interaction with nsp7 protein (yellow circle) of SARS-CoV-2.

**S2 Fig. Functional hub derived from viral replication machinery network**. This cluster is termed as hub 2 that represents the closed interconnected proteins (rectangular box) showing interaction with structural proteins nucleocaspid and non- structural proteins, nsp12, 8 and 13 (indicated in a yellow circle) of SARS-CoV-2

**S3 Fig. Functional hub derived from viral replication machinery network**. This cluster is termed as hub 3 that represents the closed interconnected proteins (rectangular box) showing no direct connectivity with SARS-CoV2 proteins. However, these proteins involved in the host inflammatory process, protein serine/threonine kinase, DNA topoisomerase, transcription factor, transcription regulator, chaperone, and ubiquitin-specific protease activity.

**S4 Fig. Functional hub derived from viral replication machinery network**. This cluster is termed as hub 4 that represents the closed interconnected proteins (rectangular box) showing interaction with nsp9, nsp1 and orf10 (indicated in a yellow circle) of SARS-CoV-2

**S5 Fig. Functional hub derived from viral replication machinery network**. This cluster is termed as hub 5 that represents the closed interconnected proteins (rectangular box) showing interaction with nsp7, and nsp11 (indicated in a yellow circle) of SARS-CoV-2.

**S6 Fig. Functional hub derived from viral replication machinery network**. This cluster is termed as hub 6 that represents the closed interconnected proteins (rectangular box) showing interaction with nuclocapsid, nsp9, and nsp15 (indicated in a yellow circle) of SARS-CoV-2.

**S7 Fig. Functional hub derived from viral replication machinery network**. This cluster is termed as hub 7 that represents the closed interconnected proteins (rectangular box) showing interaction with nuclocapsid and nsp15 (indicated in a yellow circle) of SARS-CoV-2.

**S8 Fig. Functional hub derived from viral replication machinery network**. This cluster is termed as hub 8 that represents the closed interconnected proteins (rectangular box) showing interaction with orf9b and nsp13 (indicated in a yellow circle) of SARS-CoV-2.

**S9 Fig. Functional hub derived from viral replication machinery network**. This cluster is termed as hub 9 that represents the closed interconnected proteins (rectangular box) showing interaction with nsp15 (indicated in a yellow circle) of SARS-CoV-2.

**S10 Fig. Functional hub derived from viral replication machinery network**. This cluster is termed as hub 10 that represents the closed interconnected proteins (rectangular box) showing no direct interaction with SARS-CoV-2 proteins. However, the functional enrichment analysis of these proteins showed involvement in regulation of nucleic acid, cell communication and signal transduction.

**S11 Fig. Functional hub derived from viral replication machinery network**. This cluster is termed as hub 11 that represents the closed interconnected proteins (rectangular box) showing no direct interaction with SARS-CoV-2 proteins. Functional enrichment analysis of these proteins showed involvement in peptide metabolism, cell communication, signal transduction, cell growth and/or maintenance, transport and protein metabolism.

**S1 Table**. Hubs representing the differentially expressed genes

**S2 Table**. Hubs representing the proteins associated with metabolic diseases

**S3 Table**. Hub proteins showing the association with the lung related diseases

**S4 Table**. Hub proteins relating the other viral infection reported in literature

**S5 Table**. Hub proteins relating the biological function and/or pathways.

## Notes

### Competing Interest Statement

The authors have declared no competing interest.

## References

1. Bedford J, Enria D, Giesecke J, Heymann DL, Ihekweazu C, Kobinger G, Lane HC, Memish Z, Oh MD, Sall AA, Schuchat A, Ungchusak K, Wieler LH; WHO Strategic and Technical Advisory Group for Infectious Hazards. COVID-19: towards controlling of a pandemic. Lancet. 2020;395(10229):1015-1018. PMID: 32197103.

2. Lin L, Lu L, Cao W, Li T. Hypothesis for potential pathogenesis of SARS-CoV-2 infection-a review of immune changes in patients with viral pneumonia. Emerg Microbes Infect. 2020;9(1):727-732.PMID: 32196410.

3. World Health Organization (WHO); Coronavirus disease 2019 (COVID-19) Situation Report –87.

4. Hopman J, Allegranzi B, Mehtar S. Managing COVID-19 in Low- and Middle-Income Countries. JAMA. 2020. PMID: 32176764.

5. Fauci AS, Lane HC, Redfield RR. Covid-19 - Navigating the Uncharted. N Engl J Med. 2020;382(13):1268-1269. PMID: 32109011.

6. Baud D, Qi X, Nielsen-Saines K, Musso D, Pomar L, Favre G. Real estimates of mortality following COVID-19 infection. Lancet Infect Dis. 2020 pii:S1473-3099(20)30195-X. PMID: 32171390.

7. Hussain A, Bhowmik B, do Vale Moreira NC. COVID-19 and diabetes: Knowledge in progress. Diabetes Res Clin Pract. 2020;162:108142. PMID:32278764.

8. Wu F, Zhao S, Yu B, Chen YM, Wang W, Song et al. A new coronavirus associated with human respiratory disease in China. Nature. 2020;579(7798):265-269. PMID: 32015508

9. Wang Z, Chen Z, Zhang L, Wang X, Hao G, Zhang et al. China Hypertension Survey Investigators. Status of Hyperten-sion in China: Results From the China Hypertension Survey, 2012-2015. Circulation. 2018;137(22):2344-2356. PMID: 29449338.

10. European Centre for Disease Prevention and Control. Novel coronavirus disease (COVID-19) pandemic: increased transmission in the EU/EEA and the UK – sixth update. Stockholm: ECDC; 2020.

11. Chan JF, Kok KH, Zhu Z, Chu H, To KK, Yuan S, Yuen KY. Genomic characterization of the 2019 novel human-pathogenic coronavirus isolated from a patient with atypical pneumonia after visiting Wuhan. Emerg Microbes Infect. 2020;9(1):221-236.PMID: 31987001.

12. Mehta P, McAuley DF, Brown M, Sanchez E, Tattersall RS, Manson JJ; HLH Across Speciality Collaboration, UK. COVID-19: consider cytokine storm syndromes and immunosuppression. Lancet. 2020;395(10229):1033-1034.PMID: 32192578.

13. Zhou Y, Hou Y, Shen J, Huang Y, Martin W, Cheng F. Network-based drug repurposing for novel coronavirus 2019-nCoV/SARS-CoV-2. Cell Discov. 2020;6:14. PMID: 32194980.

14. Gordon DE, Jang MG, Bouhaddou M, et al. (2020) A SARS-CoV-2-Human Protein-Protein Interaction Map Reveals Drug Targets and Potential Drug-Repurposing. BioRxiv doi: https://doi.org/10.1101/2020.03.22.002386 [PREPRINT].

15. Bader GD, Hogue CW. An automated method for finding molecular complexes in large protein interaction networks. BMC Bioinformatics. 2003;4:2. PMID: 12525261.

16. Conti P, Gallenga CE, TetÈ G, Caraffa A, Ronconi G, Younes A, Toniato E, Ross R, Kritas SK. How to reduce the likelihood of coronavirus-19 (CoV-19 or SARS-CoV-2) infection and lung inflammation mediated by IL-1. J Biol Regul Homeost Agents. 2020;34(2). PMID: 32228825.

17. Kong WH, Li Y, Peng MW, Kong DG, Yang XB, Wang et al. SARS-CoV-2 detection in patients with influenza-like illness. Nat Microbiol. 2020. PMID: 32265517.

18. Gupta P, Bhattacharjee S, Sharma AR, Sharma G, Lee SS, Chakraborty C. miRNAs in Alzheimer Disease - A Therapeutic Perspective. Curr Alzheimer Res. 2017;14(11):1198-1206.PMID: 28847283.

19. Rupaimoole R, Slack FJ. MicroRNA therapeutics: towards a new era for the management of cancer and other diseases. Nat Rev Drug Discov. 2017;16(3):203-222.PMID: 28209991.

