## Supplementary figures and images for "Interplay of host regulatory network on SARS-CoV-2 binding and replication machinery"

### S1 Fig

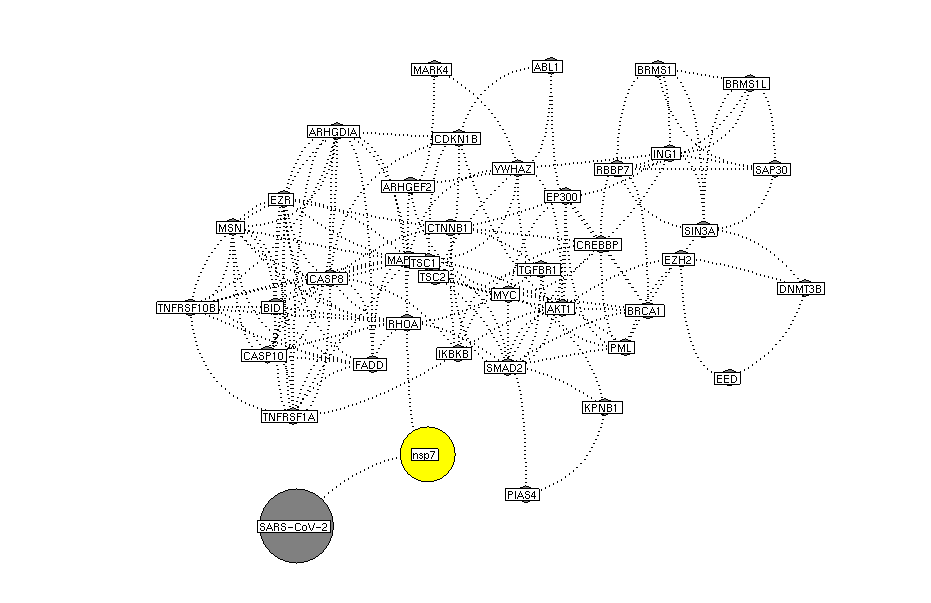

### S2 Fig

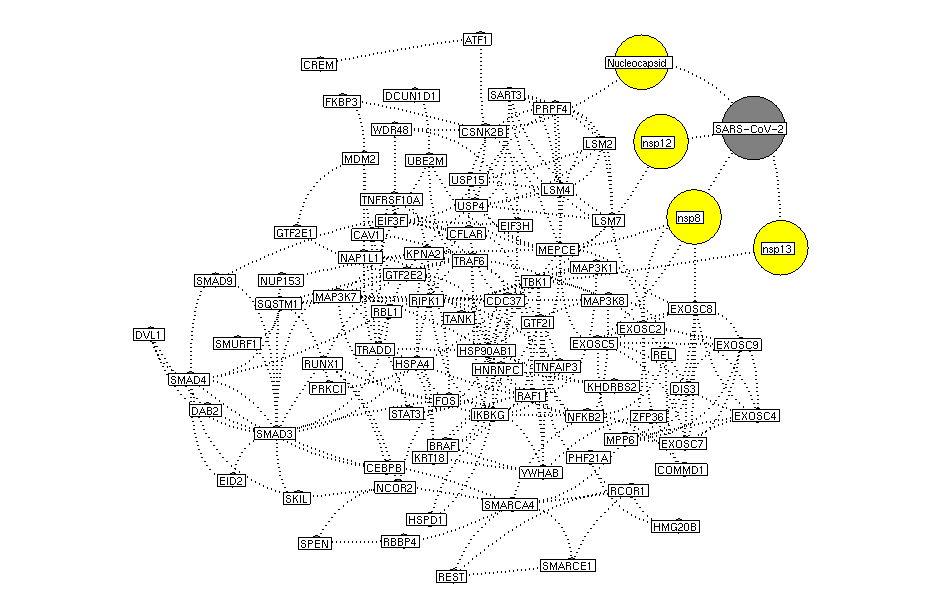

### S3 Fig

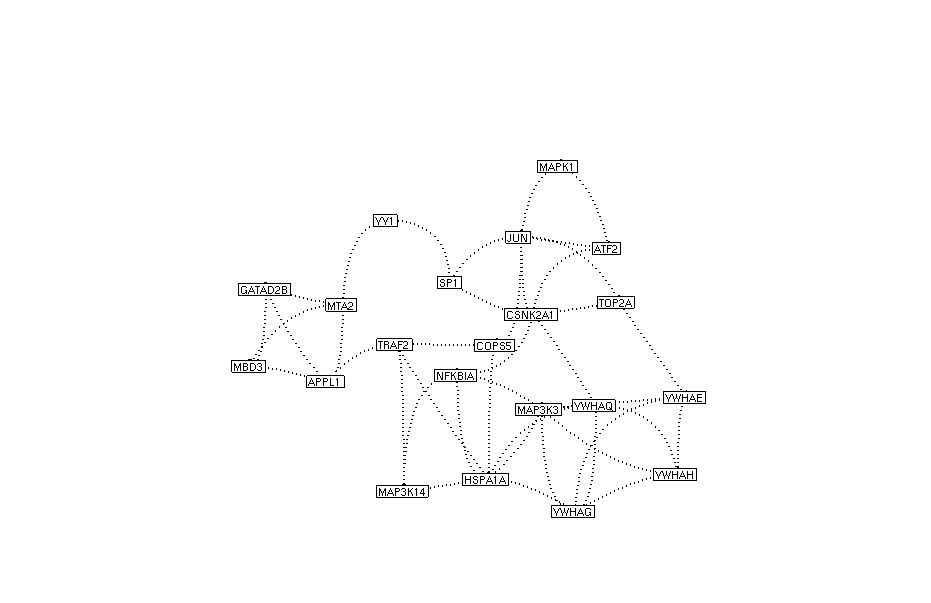

### S4 Fig

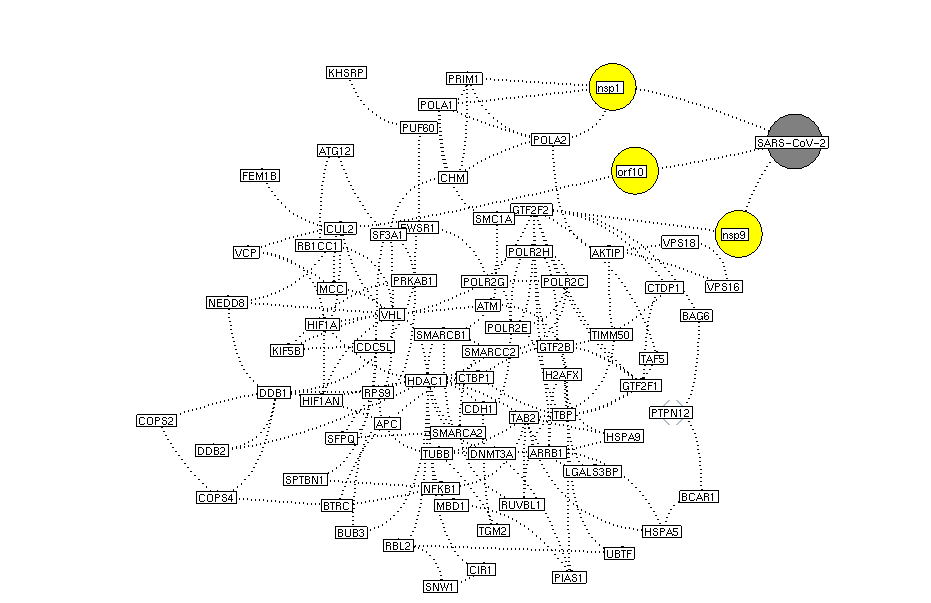

### S5 Fig

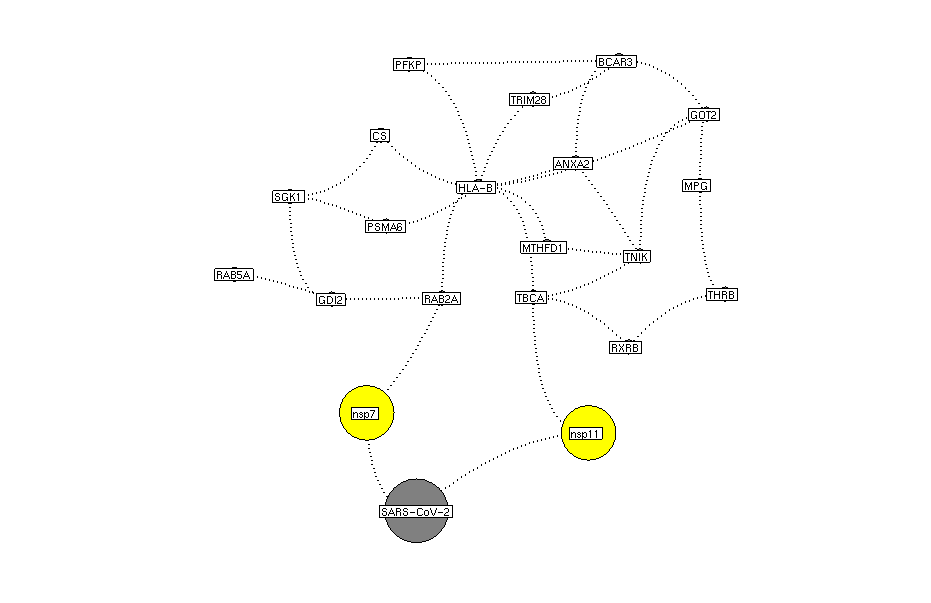

### S6 Fig

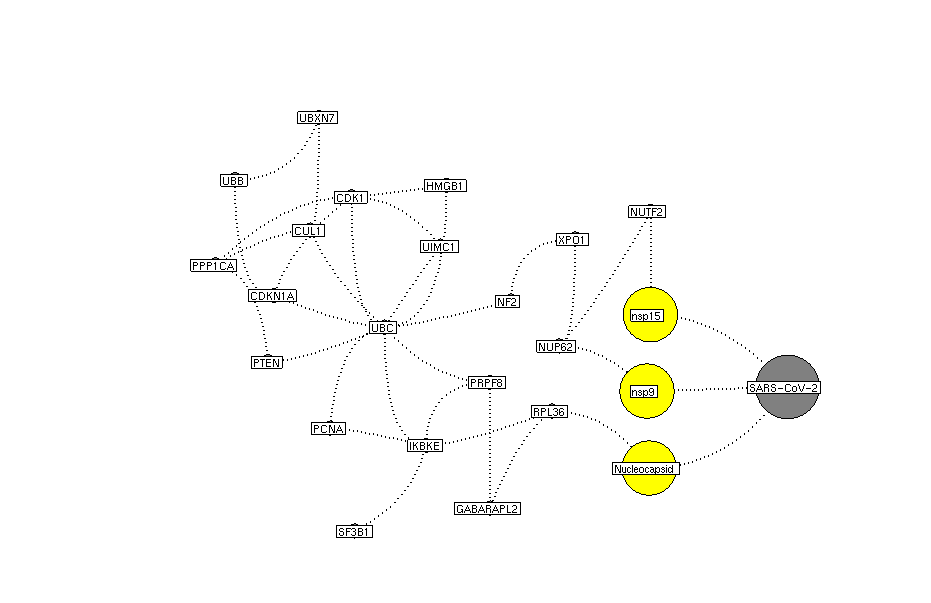

### S7 Fig

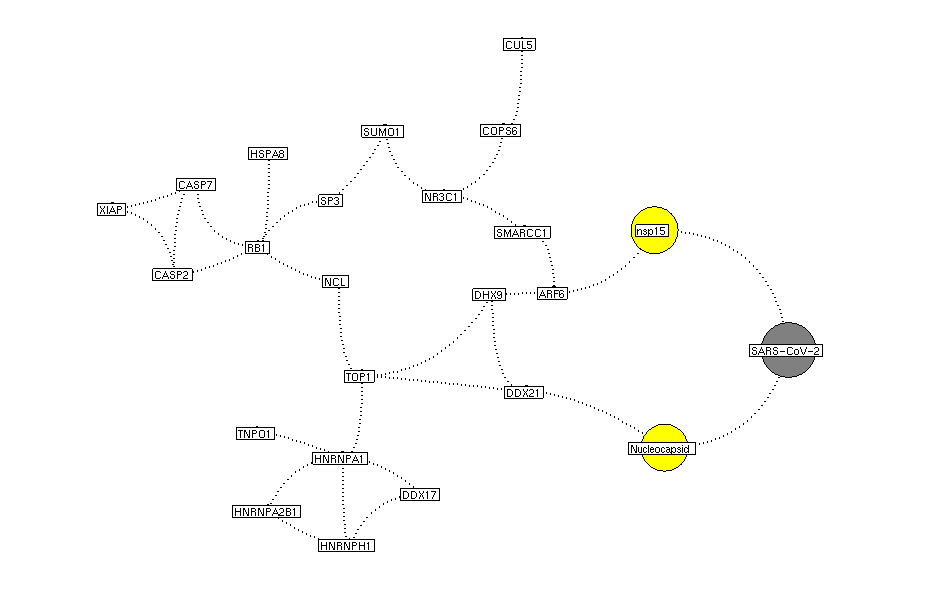

### S8 Fig

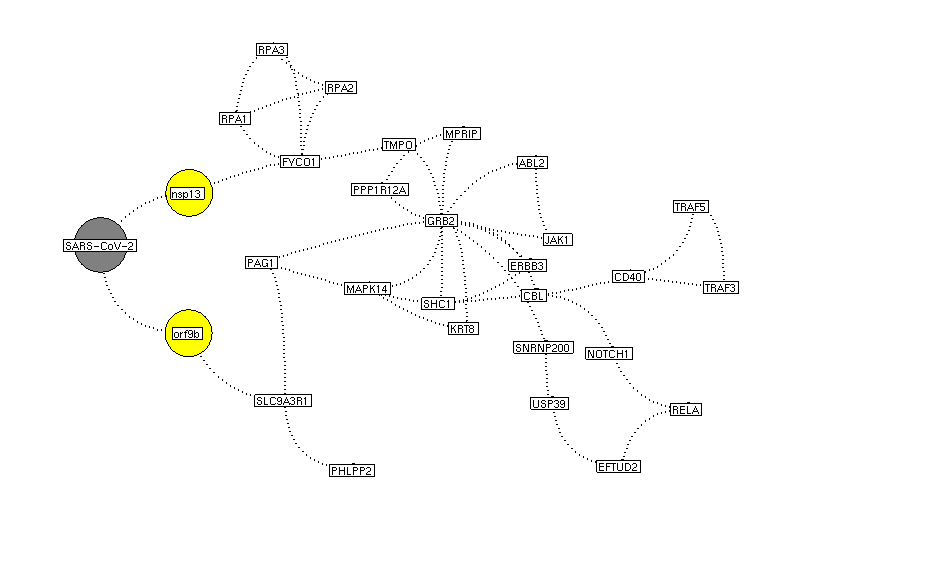

### S9 Fig

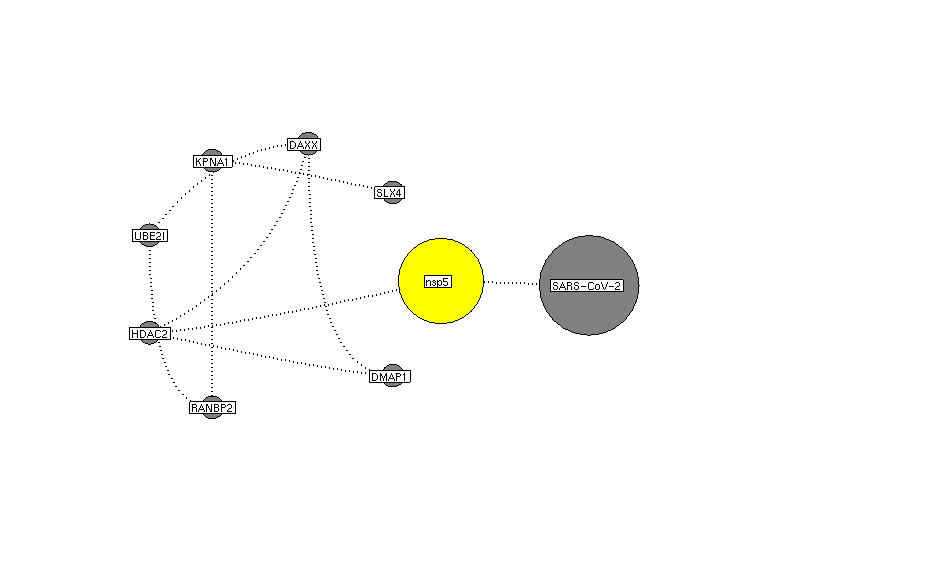

### S10 Fig

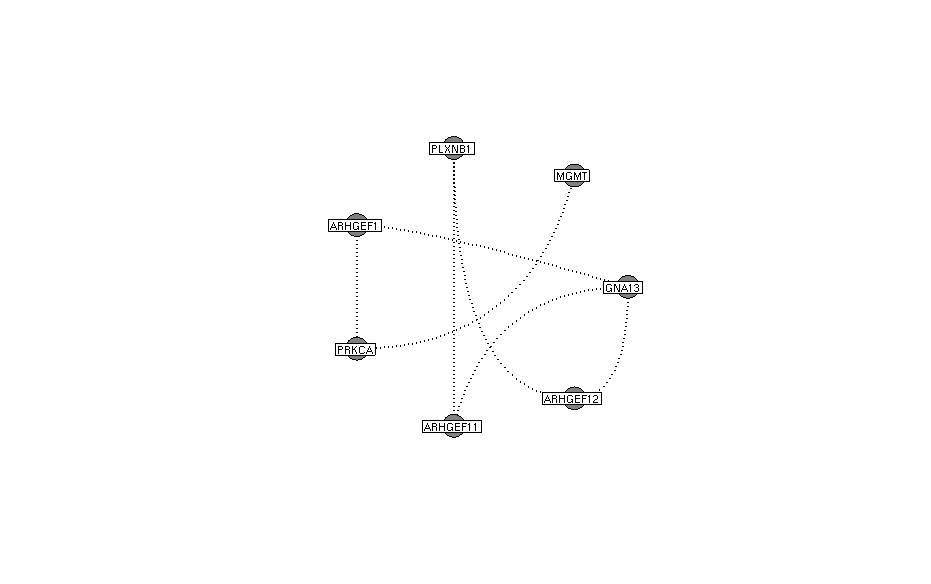

### S11 Fig

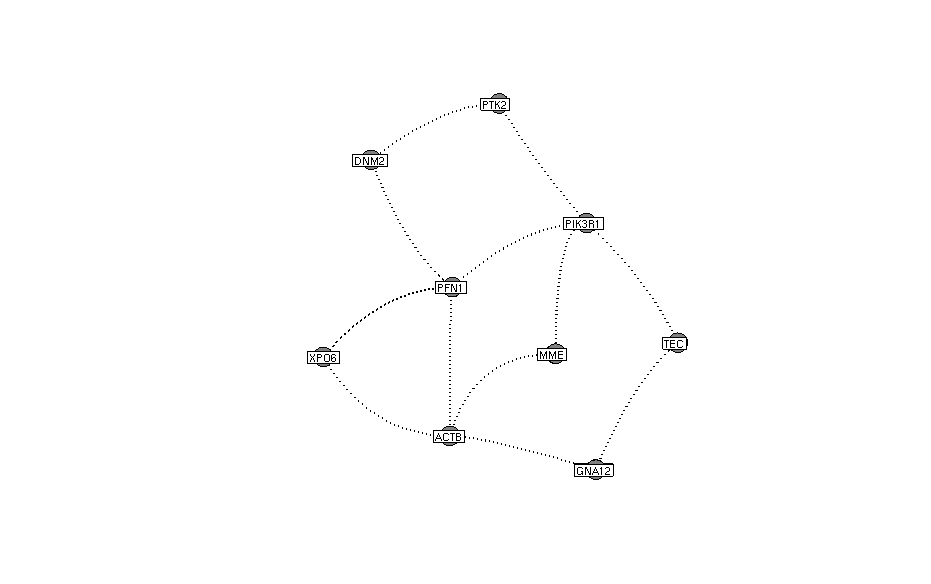
